# Aligned multicompartment collagen scaffolds support stratified myoblast and fibroblast behavior for musculotendinous tissue engineering

**DOI:** 10.64898/2026.06.12.730640

**Authors:** Geshani C. Bandara, Ryann D. Boudreau, William Wyatt, Steven R. Caliari

## Abstract

Injuries to musculoskeletal tissue junctions are exceedingly common and notoriously difficult to repair due to the inability to restore overlapping gradations of structural, biochemical, and mechanical signals critical to tissue interfacial integrity. This work introduces a multicompartment scaffold for muscle-tendon junction (MTJ) tissue engineering, containing distinct ‘muscle’ and ‘tendon’ compartments joined at a continuous interface, recapitulating the structural anisotropy, graded collagen content, and electrical excitability of the native MTJ. Collagen suspensions with or without electrically conductive poly(3,4-ethylenedioxythiophene) (PEDOT) particles representing ‘muscle’ and ‘tendon’ compartments respectively were carefully layered and directionally freeze-dried to form an integrated multicompartment scaffold with aligned pores mimicking the MTJ. Scanning electron microscopy (SEM) and energy-dispersive X-ray spectroscopy (EDS) confirmed the formation of a structurally anisotropic scaffold with stratified conductive polymer content, and importantly, a smooth continuous interfacial region joining the two compartments of similar scale to native MTJ. In contrast to multicompartment materials with abrupt interfaces, mechanical testing confirmed no decrease in multicompartment scaffold tensile properties relative to single compartment controls. Myoblasts and fibroblasts were successfully seeded on multicompartment scaffolds in a stratified manner while uniformly conforming to aligned scaffold contact guidance cues and maintaining metabolic activity over a week in culture. Myoblasts underwent compartment-specific differentiation while fibroblasts remained viable, even under myogenic differentiation conditions. Together, this work presents a scaffold platform integrating key structural, biochemical, and mechanical features necessary for MTJ tissue engineering.

## 1. Introduction

While biomaterials approaches to repair individual tissues are widely studied, the engineering of complex tissue interfaces remains relatively underexplored, despite the significant clinical need^[1]^. Multi-tissue interfaces such as cartilage-bone, meniscus-bone, tendon/ligament-bone, and muscle-tendon are ubiquitous in the musculoskeletal system and are also common injury sites despite the presence of elegant overlapping biophysical and biochemical gradations between neighboring tissues^[1–5]^. As a primary example, the musculotendinous junction (MTJ) is a critical interface within the musculoskeletal system, enabling normal locomotion via efficient force transmission between muscle-tendon-bone units^[1]^. However the MTJ, like other musculoskeletal tissue interfaces, is susceptible to injury due to the integration of mechanically and biochemically distinct tissues, which leads to localized stress concentrations during loading^[6,7]^. Consequently, MTJ injuries commonly arise under conditions such as excessive stretching, high-impact physical activity, and aging^[8–10]^. Moreover, many skeletal muscle injuries involve MTJ damage, particularly in severe cases such as volumetric muscle loss (VML) injuries resulting from traumatic events including vehicle accidents and combat-related injuries^[11–13]^.

The native MTJ is a highly specialized interface composed of three distinct yet functionally integrated regions: 1) muscle, an elastic and an electrically responsive tissue consisting of aligned multinucleated muscle fibers encased in collagen and laminin-rich connective extracellular matrix (ECM), 2) tendon, a stiff, collagen-rich tissue with sparsely distributed fibroblasts, and 3) the MTJ, where electrically responsive and excitable muscle fibers and sarcolemma (muscle cell membrane) integrate with the collagen-dense tendon ECM^[1,14–19]^. This hierarchical architecture enables efficient transmission of contractile forces generated by muscle to the tendon, facilitating coordinated movement and locomotion^[20,21]^. Notably, as in other musculoskeletal junctions like the tendon-bone junction^[22]^, the transition zone between muscle and tendon is quite small, with widths on the order of tens to hundreds of microns^[8,15,23–25]^. Despite these structural and functional insights, engineering biomaterial-based MTJ scaffolds remains challenging due to the need to recapitulate the mechanical, biochemical, electrical, and cellular gradients present in native tissue.

Previous efforts to engineer biomaterial-based MTJ constructs have largely focused on recreating the graded mechanical environment of the native interface through compartmentalized scaffold designs and spatially controlled mechanical properties. A variety of fabrication strategies, including electrospinning^[26]^, bioprinting^[27]^, and artificial intelligence-assisted 3D printing^[28]^ have been employed to generate mechanically distinct muscle- and tendon-like regions capable of supporting tissue-specific cellular responses and MTJ-associated gene expression. Collectively, these studies demonstrate that incorporating stiffness gradients enhances physiological relevance of engineered MTJ models and promotes the development of interface-specific phenotypes. Beyond mechanical design, recent advances have increasingly incorporated tissue-specific biochemical cues to regulate cell behavior and muscle and tendon regeneration. The inclusion of tissue-specific ECM components^[17,29]^ and cellular signals^[30]^ has been shown to modulate cell differentiation and, matrix deposition and tissue maturation within MTJ constructs. While these findings separately highlight the importance of biochemical and mechanical cues for engineering MTJs, there remains a need to integrate graded biomimetic biochemical and mechanical cues into a single scaffold construct.

In addition to biochemical and mechanical signals, the role of electrical responsiveness in MTJ scaffolds remains largely unexplored, representing a critical gap in the design of biomimetic MTJ constructs. Electrical signals play a crucial role in regulating cellular processes within electrically-responsive tissues such as skeletal muscle, influencing cell proliferation and differentiation^[31–36]^. Conductive biomaterials have been shown to enhance multinucleated myotube formation *in vitro* and promote myofiber regeneration *in vivo* compared to non-conductive biomaterials by our group^[37,38]^ and others^[39]^, underscoring the importance of electrical cues in muscle tissue regeneration. Accordingly, incorporating electrical functionality into MTJ constructs may provide important biophysical cues that complement the structural, mechanical, and biochemical features of the native interface. Conductive polymers, including polypyrrole (PPy), poly(3,4-ethylenedioxythiophene) (PEDOT), and polyaniline (PANi) have emerged as promising materials for imparting electrical conductivity to tissue-engineered scaffolds due to their tunable electrical properties and demonstrated ability to support muscle cell function and maturation^[36,40,41]^.

Building upon these insights, the present study integrates both the anisotropic structural organization and electrical responsiveness gradient of the native MTJ into a single biomaterial by engineering an aligned, multicompartment collagen scaffold via directional freeze-drying. The resulting scaffold comprises stratified electrically-conductive ‘muscle’ and non-conductive ‘tendon’ compartments connected through a continuous interface of similar scale to native MTJs. Using this platform we investigate compartment-specific cellular alignment, metabolic activity, and differentiation with the goal of developing a scaffold platform for MTJ tissue engineering.

## 2. Materials and Methods

### 2.1 Glycosaminoglycan (GAG)-doped poly(3,4-ethylenedioxythiophene) synthesis

Glycosaminoglycan (GAG)-doped PEDOT particles were synthesized by initially mixing 1.6 mL (0.015 mol) of EDOT (Sigma Aldrich) monomer in 50 mL of deionized (DI) water at 30°C. In a separate flask, 5.13 g (0.02 mol) of ammonium persulfate (APS, Sigma Aldrich) and 75 mg of GAG (hyaluronic acid (HA), 19 kDa, Lifecore or heparin (HP), 5-15 kDa, Sigma Aldrich) were dissolved in 50 mL of DI water. The APS-GAG solution was then added dropwise to the EDOT solution under vigorous stirring. The reaction proceeded at 30°C for 24 hours. The resulting black precipitate was separated via centrifugation at 5000 rpm for 5 min by adding an equal volume of acetone. Further purification was performed through sequential centrifugation steps; once with ethanol and three times with DI water to remove impurities. The collected precipitate was then dried in a vacuum oven at 60°C overnight. Finally, the dried particles were passed through a 325-mesh (45 μm) screen to obtain the final GAG-doped PEDOT particles.

### 2.2 Scaffold fabrication

Scaffolds were fabricated through directional freeze-drying of collagen-GAG (CG) suspension as described in our previous papers (**Figure 1**)^[37,42,43]^. 1.5 wt/v% or 2.5 wt/v% collagen suspensions were made by homogenizing microfibrillar type I collagen (bovine Achilles tendon) in 0.05 M acetic acid solution for conductive or non-conductive scaffold compartments respectively. The suspensions were homogenized at 15000 rpm in a recirculating chiller maintained at 4°C to avoid collagen denaturation caused by shear-induced heating. A precooled heparin solution in 0.05 M acetic acid was added dropwise to achieve a final 0.1 wt/v% GAG concentration. For conductive scaffolds, 1 wt/v% GAG-doped PEDOT particles were incorporated into the homogenized CG suspension by vortexing before freeze-drying. Single compartment scaffolds (conductive and non-conductive) were fabricated by pipetting respective CG suspensions with and without conductive polymer particles into a thermally-mismatched Teflon mold with a copper base, facilitating longitudinal heat transfer. Multicompartment scaffolds were produced by carefully layering the non-conductive suspension on top of the conductive suspension and allowing interdiffusion for 30 min prior to freeze-drying^[44]^. The filled molds were then placed on a pre-cooled −10°C shelf and directionally freeze-dried in a VirTis Genesis pilot scale freeze-dryer. After freeze-drying, scaffolds were placed in a vacuum oven at 105 °C for 24 h to promote dehydrothermal crosslinking to improve scaffold stability and mechanical properties^[37]^. Scaffolds were then cut into cylindrical discs with a diameter of 6 mm and a height of 4-5 mm. Multicompartment scaffolds were sectioned such that the interface was located at the center of each scaffold.

**Figure 1.**
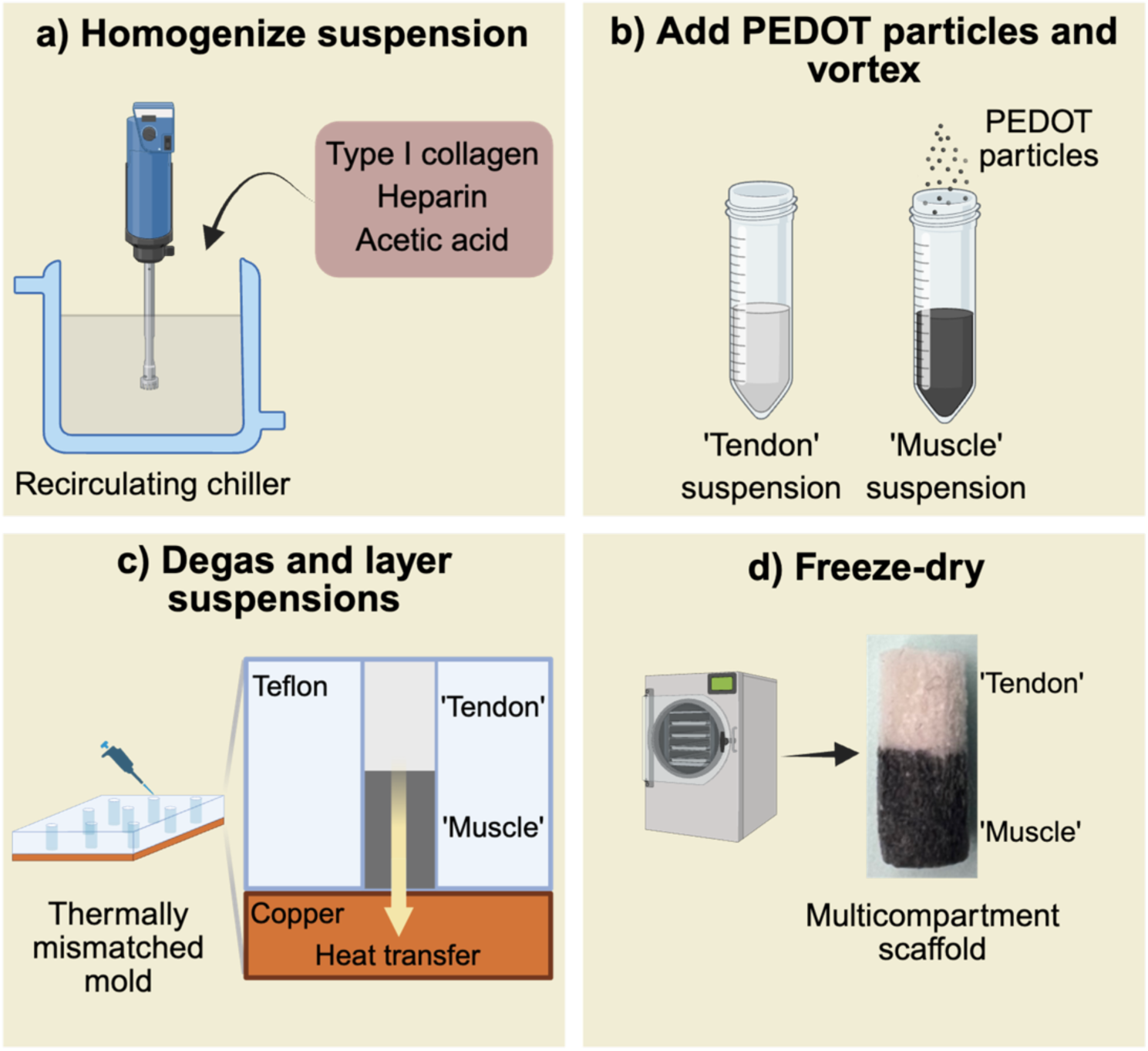
Schematic of the multicompartment scaffold fabrication approach. a) Suspensions of type I collagen and heparin in acetic acid are homogenized to start the process. b) To make the ‘muscle’ compartment electrically conductive, GAG-doped PEDOT particles are mixed into the collagen-heparin suspension. c) Conductive and non-conductive precursor suspensions are then sequentially layered to generate distinct regions mimicking ‘muscle’ (conductive) and ‘tendon’ (non-conductive) compartments. d) The assembled mold is subjected to directional freeze-drying by placing on a pre-cooled freeze-dryer shelf at −10 °C. Controlled directional heat transfer during freezing promotes formation of an aligned porous architecture mimicking the MTJ. Figure adapted and modified from Basurto et. al^[37]^.

### 2.3 Scanning electron microscopy and energy-dispersive X-ray spectroscopy analysis

Scanning electron microscopy (SEM) was used to characterize the morphology of conductive particles and the pore architecture of dry scaffolds. Particle imaging was performed using a secondary electron detector under high vacuum conditions (0.1 Pa), while scaffold imaging was conducted using a backscattered electron detector under low vacuum conditions (60 Pa) on a Phenom ESEM operated at an accelerating voltage of 10 kV. The diameters of 150 conductive particles were measured across three distinct fields of view using the ‘Measure’ function in ImageJ software to evaluate particle size distribution. Pore orientation was quantified using the OrientationJ plugin in ImageJ. Energy-dispersive X-ray spectroscopy (EDS) mapping was performed in conjunction with SEM to visualize the spatial distribution of conductive particles within the scaffolds. The presence of sulfur atoms, originating from the oxidizing agent ammonium persulfate (APS) and the PEDOT backbone, was used to map localization of GAG-doped PEDOT particles. PEDOT particle distribution was quantified by analyzing the spatial intensity distribution of sulfur-containing pixels from EDS maps using ImageJ.

### 2.4 Scaffold hydration, crosslinking, fluorescent labeling, and sterilizing

Scaffolds were hydrated by soaking in 70% ethanol for 20 min, followed by two rinses with phosphate-buffered saline (PBS). Subsequently, scaffolds were immersed in a 2 μM solution of AlexaFluor 568 NHS ester in PBS for 20 min to label the primary amines in the collagen backbone. Chemical crosslinking was then performed by gently shaking the scaffolds in a solution of 1-ethyl-3-(3-dimethylaminopropyl) carbodiimide hydrochloride (EDC) and N-hydroxysulfosuccinimide (NHS) solution at a molar ratio of 5:2:1 (EDC:NHS:COOH), where COOH represents the carboxylic acid content of collagen^[46]^. Finally, the scaffolds were immersed in 70% ethanol for 2 h and in sterilized PBS overnight prior to cell seeding.

### 2.5 Scaffold conductivity measurements

Hydrated and crosslinked scaffolds were used for all measurements. PBS was substituted with deionized water in every step to minimize ionic interference in measurements. Before conductivity measurements, scaffold height and diameter were measured. Conductivity was assessed using a parallel plate cell with copper electrodes, connected to a Biologic Science Instrument controlled via EC-Lab® software. The scaffolds were placed between the parallel plates, ensuring uniform contact by adjusting the electrode spacing to match the scaffold height. Linear sweep voltammetry (LSV) was performed by sweeping the voltage from −1V to 1V, while recording the resulting current. A current versus voltage graph was obtained, where the slope of the linear region of the graph represented the reciprocal of the resistance (R). The electrical conductivity (δ) was calculated by Pouillet’s law (**Equation 1**) where L is the length of the scaffold and A is the cross-sectional area of the scaffold in contact with the electrodes (A = πD^2^/4, D is the scaffold diameter). Three scaffolds per experimental group were tested.

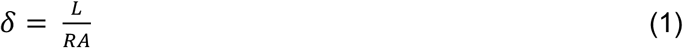

### 2.6 Scaffold mechanical characterization

Scaffold mechanical properties (single compartment ‘muscle’ (M), ‘tendon’ (T), or multicompartment ‘muscle-tendon’ (M-T) scaffolds) were assessed using tensile testing in the dry state. Scaffolds were fabricated in a larger mold (1 cm diameter and 3 cm height) and subsequently cut into a dog-bone geometry using a razor blade. To prevent local compression near the grips during testing, the scaffold ends were reinforced by gluing and covering with cardboard strips, enabling secure clamping. Tensile testing was performed using an Instron equipped with a 5 N load cell, and samples were strained until failure. The width and the thickness of each sample were measured using calipers prior to testing. Stress-strain curves were generated by force and displacement values acquired using Bluehill software, and the Young’s modulus was calculated from the slope of the linear regions of the stress-strain curves. Ultimate tensile strength was determined as the maximum stress value corresponding to the highest point of the stress-strain curve prior to failure.

### 2.7 Scaffold pore size analysis

Hydrated, crosslinked, and fluorescently-labeled scaffolds were imaged using a Cytation C10 confocal microscope to analyze pore morphology. Images with a 10 μm z-thickness were selected for analysis. Pores were segmented using Omnipose, followed by manual correction to refine the segmentation. The lengths of pore major axis (a) and minor axis (b) were calculated using the region_props function from the scikit-image Python package. The pore diameter (d) was then calculated by fitting each pore to an ellipse using **Equation 2**.

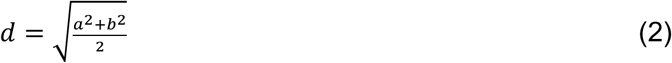

### 2.8 Cell culture

Immortalized mouse myoblasts (C2C12, ATCC) and fibroblasts (NIH 3T3, Sigma) at passages 2-4 were cultured in standard T75 tissue culture flasks using growth media consisting of high-glucose Dulbecco’s Modified Eagle’s Medium (DMEM, ThermoFisher) supplemented with 10 v/v% fetal bovine serum (FBS, Gibco) and 1 v/v% antibiotic-antimycotic solution (Invitrogen). Cells were passaged or used in experiments upon reaching 80% confluence. Myogenic differentiation was induced using differentiation media composed of DMEM supplemented with 2 v/v% horse serum (Gibco) and 1 v/v% antibiotic-antimycotic. All cell cultures were maintained at 37 °C in a humidified incubator with 5% CO_2_, and media were refreshed every other day.

### 2.9 Scaffold culture conditions

Hydrated and crosslinked scaffolds were rinsed with sterile PBS and preconditioned in growth media for 30 min prior to cell seeding. Cell suspensions were prepared by trypzinizing confluent cultures and resuspending cells at a concentration of 2×10^5^ cells per 40 μL. Scaffolds were gently blotted on sterile wipes to remove excess media and transferred into ultra-low attachment six-well plates to minimize non-specific cell adhesion. To establish compartment-specific cellular organization, 10 μL aliquots of C2C12 myoblast and NIH 3T3 fibroblast suspensions were pipetted onto the central and lateral regions of the M and T compartments respectively. At the M-T interface region, 5 μL of each cell suspension was co-seeded to promote formation of a mixed interfacial cell population. Following an initial 20 min incubation to facilitate cell attachment, scaffolds were inverted and the seeding procedure was on the opposite surface, excluding the lateral regions. Scaffolds were subsequently incubated for 2 h prior to the addition of growth media. Growth media was replaced after 24 h and subsequently every other day. On day 4, scaffolds were transitioned to differentiation media, which was similarly refreshed every other day for the remainder of the culture period. All cultures were maintained under standard incubation conditions at 37 °C and 5% CO_2_.

### 2.10 Cell metabolic activity

Cell metabolic activity was evaluated using the alamarBlue assay. To enable compartment-specific analysis, scaffolds were dissected at the M-T region immediately prior to the assay and the resulting M and T regions were incubated separately in 10% alamarBlue solution for 70 min on a shaker at 80 rpm. Metabolic activity was quantified by measuring the fluorescence intensity of fluorescent resorufin, the fluorescent pink reduction product generated from the conversion of non-fluorescent blue resazurin present in the alamarBlue solution through mitochondrial and cytoplasmic redox reactions of viable cells^[47]^. Fluorescence measurements were acquired using a plate reader (Tecan M200, excitation: 565 nm, emission: 595 nm).

### 2.11 Fluorescent labeling, Immunocytochemistry and confocal imaging

For cell tracking studies, C2C12 myoblasts and NIH 3T3 fibroblasts were fluorescently labeled prior to seeding using green fluorophore DiO (Vybrant® DiO Cell-Labeling Solution, Thermo Fisher Scientific) and red fluorophore DiI (Vybrant® DiI Cell-Labeling Solution, Thermo Fisher Scientific), respectively. Briefly, 5 µL of fluorophore solution was added to cell suspension at a density of 1×10^5^ cells/mL prepared in serum-free media and incubated for 20 min at 37 °C. Following labeling, cells were centrifuged, washed twice with growth media, and subsequently seeded into scaffolds. At the conclusion of the culture period scaffolds were fixed in 10% neutral-buffered formalin, rinsed with PBS, and prepared for confocal imaging.

For immunocytochemical analysis, fixed scaffolds were permeabilized in 0.1% Triton X-100 in PBS for 30 mins, followed by blocking with 3 % (w/v) BSA in PBS for 1 h to minimize non-specific antibody binding. To assess myogenic differentiation, scaffolds were incubated overnight at 4 °C with a primary anti-myosin heavy chain (MHC) antibody (Myosin 4 monoclonal antibody (MF20), eBioscience) diluted 1:400 in 3% BSA. Scaffolds were subsequently rinsed three times with PBS and then incubated for 2 h at room temperature with AlexaFluor 488-conjugated goat anti-mouse IgG (1:400 dilution in 3% BSA; Thermo Fisher Scientific) as the secondary antibody. Simultaneously, fluorescein phalloidin (1:400 dilution in 3% BSA; Thermo Fisher Scientific) was used to visualize F-actin and cell cytoskeletal organization. Following secondary staining, scaffolds were rinsed three times with 0.05% Tween in PBS and incubated with DAPI (4′,6-diamidino-2-phenylindole, Thermo Fisher Scientific) for 5 min to visualize cell nuclei. Scaffolds were then rinsed twice with PBS and stored in the dark at 4 °C prior to confocal microscopy imaging.

Confocal imaging was performed using a Cytation 10 confocal microscope. Maximum projections of z-stack images acquired over a thickness 100 μm were generated from three independent fields of view per scaffold region and were used for quantitative analysis. For scaffold pore size quantification, single sections corresponding to the scaffold backbone were extracted from the z-stack images. The scaffold backbone, F-actin cytoskeleton, MHC expression, and cell nuclei were visualized using TRITC, CY5, GFP, and DAPI fluorescence channels, respectively. Image segmentation was performed using Omnipose, incorporating both nuclei and MHC channels followed by a manual correction to ensure segmentation accuracy. Quantitative cellular morphological analysis was subsequently performed using the region_props function implemented in the scikit-image Python package^[48,49]^.

### 2.12 Statistical analysis

Statistical analyses were performed using one-way or two-way ANOVA followed by Tukey’s HSD post hoc test, or Welch’s t-test where appropriate, using GraphPad Prism (Version 11). Statistical significances are defined as p < 0.05, 0.01, 0.001, or 0.0001, with significance levels denoted as ∗, ∗∗, ∗∗∗, and ∗∗∗∗, respectively. Data are represented as mean ± standard deviation. Bar graphs and violin plots include overlaid scatter points representing individual biological replicates. Each experiment was performed using three or four scaffolds as biological replicates. For immunocytochemical analyses, three regions of interest (ROIs) were analyzed per scaffold region and averaged for statistical analysis.

## 3. Results and Discussion

### 3.1 Scaffolds incorporating GAG-doped PEDOT particles support myoblast metabolic activity and myogenic differentiation

The incorporation of conductive polymers into biomaterial scaffolds has become a promising strategy in numerous tissue engineering applications to not only provide structural support for cellular attachment but also to deliver biomimetic electrical cues that modulate cellular function^[40]^. Despite these advantages, the integration of conductive polymers into tissue engineering scaffolds remains challenging due to limitations such as nonbiodegradability, potential cytotoxicity, and difficulty in achieving uniform distribution within biomaterial matrices^[32]^.

In our previous work, CG scaffolds incorporating HA-doped PEDOT particles supported significantly enhanced myoblast metabolic activity compared to scaffolds with equivalent electrical conductivity incorporating PPy particles, highlighting the superior cytocompatibility of PEDOT over PPy within CG scaffolds^[38]^. We also recently showed that non-conductive CG scaffolds incorporating HP promoted superior myogenic differentiation and growth factor sequestration^[43]^. This motivated us to directly compare the performance of conductive scaffolds incorporating PEDOT particles doped with either HA or HP as a preliminary step to fabricating multicompartment scaffolds.

PEDOT-HA and PEDOT-HP particles of similar sizes (∼ 190-230 nm diameter) were successfully synthesized and homogenously incorporated into CG scaffolds (**Figure S1a-g**). Incorporation of both PEDOT-HA and PEDOT-HP particles significantly increased scaffold bulk conductivity relative to collagen scaffold controls without conductive particles (**Figure S1h**). C2C12 mouse myoblasts grown in both conductive scaffold groups showed similar metabolic activity (**Figure S2**) and myogenic differentiation capacity (**Figure S3**) compared to non-conductive controls. Collectively, these findings demonstrate the successful fabrication of conductive GAG-doped PEDOT scaffolds capable of introducing enhanced electrical functionality while maintaining a milieu suitable for musculoskeletal tissue engineering. While results between the PEDOT-HA and PEDOT-HP groups were similar, we chose to move forward with PEDOT-HA as our conductive polymer in the ‘muscle’ (M) compartment due to our use of it in previous work^[38]^ as well as the slightly higher metabolic activity and myogenic differentiation levels compared to the PEDOT-HP group.

### 3.2 Multicompartment scaffolds show graded conductivity with an interface comparable to native MTJ scales

Multicompartment scaffolds were fabricated as depicted in **Figure 1** by layering ‘tendon’ (T) suspension (2.5 wt/v% collagen, 0.1 wt/v% HP) on top of ‘muscle’ (M) suspension (1.5 wt/v% collagen, 0.1 wt/v% heparin, 1 wt/v% PEDOT-HA) and freeze-drying. We hypothesized that careful suspension layering and interdiffusion prior to freeze-drying would result in a smooth continuous interface between the M and T compartments reminiscent of native musculoskeletal junctions like the MTJ. We first characterized compartment integration by assessing the spatial distribution of PEDOT within the scaffold using EDS elemental mapping of sulfur in conjunction with SEM. Sulfur signals corresponding to PEDOT were overlaid to SEM images to visualize particle localization across scaffold regions. The M compartment exhibited a dense and homogenous distribution of PEDOT particles, as indicated by the strong sulfur signal (**Figure 2a**). In contrast, the M-T region displayed a transition in particle density between compartments (**Figure 2b**). Weaker sulfur signal was observed in the T compartment, likely due to the presence of heparin (**Figure 2c**).

**Figure 2.**
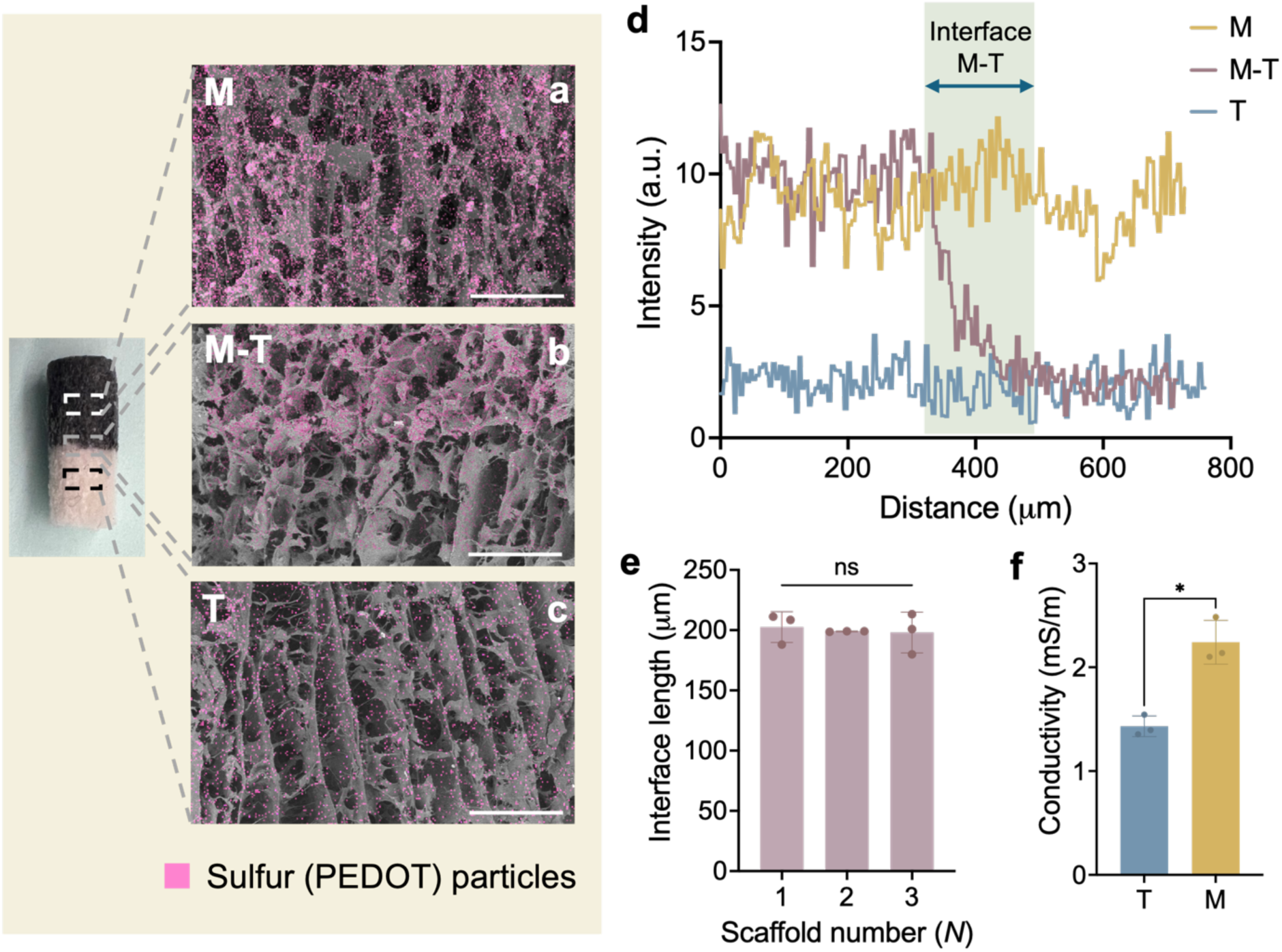
Multicompartment scaffolds show graded conductivity with an interface matching native MTJ scales. a-c) Energy-dispersive X-ray spectroscopy (EDS) sulfur maps (pink), corresponding primarily to PEDOT, overlaid on scanning electron microscopy (SEM) images of longitudinal scaffold sections, highlighting the ‘muscle’ (M), ‘muscle-tendon junction’ (M-T), and ‘tendon’ (T) regions. PEDOT particles are densely and uniformly distributed within the muscle compartment and then decrease across the interface, indicating successful compartment and PEDOT particle stratification. Scale bars: 300 µm. d) Quantitative analysis of PEDOT distribution, showing sulfur intensity as a function of distance across compartments, confirms a smooth and continuous transition at the interface. e) The resulting interface exhibits consistent and reproducible dimensions, with an average length of 200 ± 11 µm (One-way ANOVA with Tukey’s HSD post hoc tests, ns: no statistically significant differences. *N* = 3). f) Conductivity measurements demonstrate a significant increase in electrical conductivity in the M compartment compared to the T compartment (Welch’s t-test, ✽: p < 0.05. *N* = 3).

To quantify the interface width, we plotted the EDS intensity profile across sections of the M and T compartments as well as at the M-T interface (**Figure 2d**). These plots highlight a linear reduction in signal at the M-T interface, reflecting a diffusion-mediated graded transition between compartments indicative of a smooth and well-defined interface. Calculating the distance over which EDS signal intensity decreased from the bulk M levels to bulk T levels revealed an average interface width of 200 ± 11 µm, which is comparable to the characteristic length scale of the native MTJ^[8,15,23–25]^. This measurement also exhibited minimal variability across samples, highlighting the reproducibility of the fabrication process (**Figure 2e**).

Importantly, incorporation of PEDOT particles resulted in a significant increase in electrical conductivity in the M compartment compared to T compartment, confirming the establishment of a functional conductivity gradient (**Figure 2f**). Collectively, these results demonstrate that the layering assembly strategy enables formation of spatially graded electrical conductivity zones joined at a continuous interface reminiscent of the MTJ.

### 3.3 Multicompartment scaffolds exhibit longitudinally aligned pores independent of PEDOT incorporation and collagen content

After characterizing the multicompartment scaffold interface, we next characterized scaffold pore architecture in both the longitudinal and transverse planes using SEM and confocal imaging. In the longitudinal direction, all compartments exhibited highly aligned pore structures (**Figure 3a**). This anisotropic architecture arises from directional ice crystal formation during freeze-drying, driven by unidirectional heat transfer imposed by the thermally-mismatched mold. Subsequent sublimation of these aligned ice crystals yields a continuous, longitudinally oriented pore network. Quantitative analysis using polar histograms further confirmed this alignment, with a strong orientation centered around 0° (direction of heat transfer) in all regions (**Figure 3c**). In contrast, transverse planes displayed a more isotropic and randomized pore morphology across all three regions (**Figure 3b**), as reflected by the broader angular distribution in the corresponding polar plots (**Figure 3d**). Pore size analysis performed using confocal images of hydrated and fluorescently-labeled scaffolds revealed comparable pore diameters across the M, M-T, and T regions in both the longitudinal and transverse planes (**Figure 3e-f, Table S1**). Collectively, these results demonstrate that the multicompartment scaffolds maintain a continuous and uniform anisotropic pore architecture throughout the construct irrespective of PEDOT incorporation and collagen content. Importantly, the presence of PEDOT particles does not disrupt ice crystal nucleation or growth during freeze-drying, indicating that scaffold microstructure is primarily governed by freeze-drying parameters rather than conductive polymer particle distribution.

**Figure 3.**
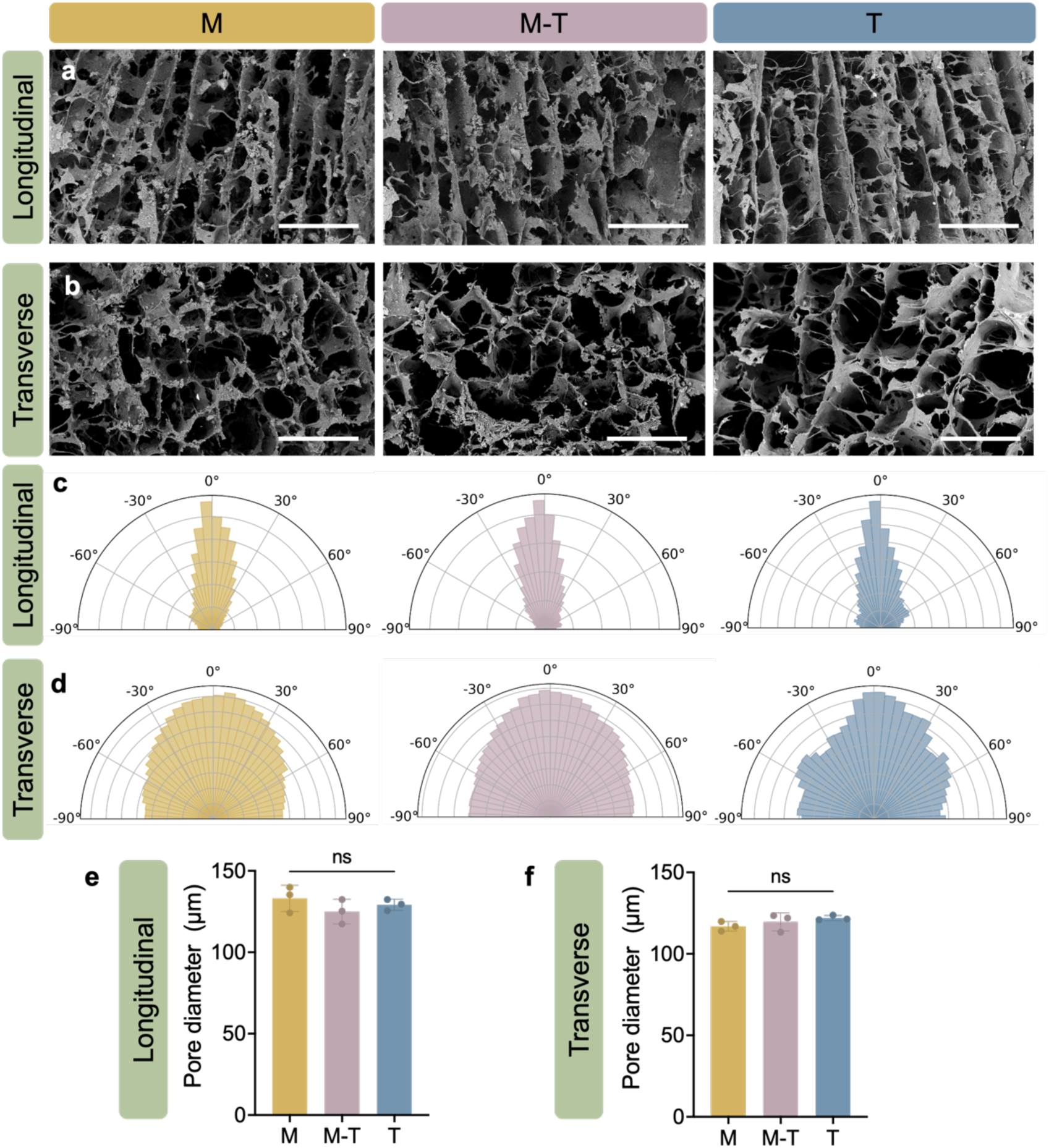
Multicompartment scaffolds exhibit longitudinally aligned pores independent of PEDOT incorporation and collagen content. a) Scanning electron microscopy (SEM) images of scaffold longitudinal plane reveal highly aligned pore structures across the ‘muscle’ (M), ‘muscle-tendon junction’ (M-T), and ‘tendon’ (T) compartments, recapitulating the architecture of native MTJ tissue. b) Corresponding transverse sections display more randomized pores in all compartments. Scale bars: 300 μm. c) Quantitative analysis of pore orientation using polar histograms demonstrates strong alignment along the longitudinal axis, d) whereas transverse sections exhibit a broader, more uniform angular distribution, confirming structural anisotropy. e) Quantification of hydrated scaffold pore diameter in the longitudinal and f) transverse planes reveals no significant differences between scaffold compartments. One-way ANOVA with Tukey’s HSD post hoc tests, ns: no statistically significant differences. *N* = 3 scaffolds per group.

### 3.4 Integrating non-conductive and conductive compartments into a continuous multicompartment scaffold does not diminish mechanics

Following characterization of multicompartment scaffold structural anisotropy, we next sought to measure scaffold mechanical integrity. Mechanical failure due to interfacial stress concentrations is a common failure mode in both musculoskeletal junction injuries and multicompartment biomaterial scaffold design. While many multicompartment scaffold designs include hard and abrupt interfaces prone to mechanical failure, we hypothesized that the diffusion-mediated interfacial gradient joining the M and T compartments would result in scaffold mechanical properties similar to single compartment scaffolds made from M or T suspensions.

We tested this hypothesis by performing uniaxial tensile testing on single compartment M and T as well as multicompartment M-T scaffolds in the dry state (**Figure 4a-c**). Representative stress-strain profiles from all scaffolds showed soft tissue-like mechanical behavior characterized by gradual elastic deformation followed by failure, with no observable mechanical discontinuities across the multicompartment construct. M-T scaffolds consistently failed within the more compliant M compartment rather than at the interface, indicating successful integration between compartments and effective load transfer across the engineered interface. The M compartment displayed greater strain to failure and increased compliance relative to the T compartment, consistent with the mechanically softer and more deformable nature of skeletal muscle tissue^[50]^. In contrast, the T compartment exhibited reduced extensibility and increased stiffness, reflecting the denser collagen network and load bearing associated with tendon-like matrices^[51]^.

**Figure 4.**
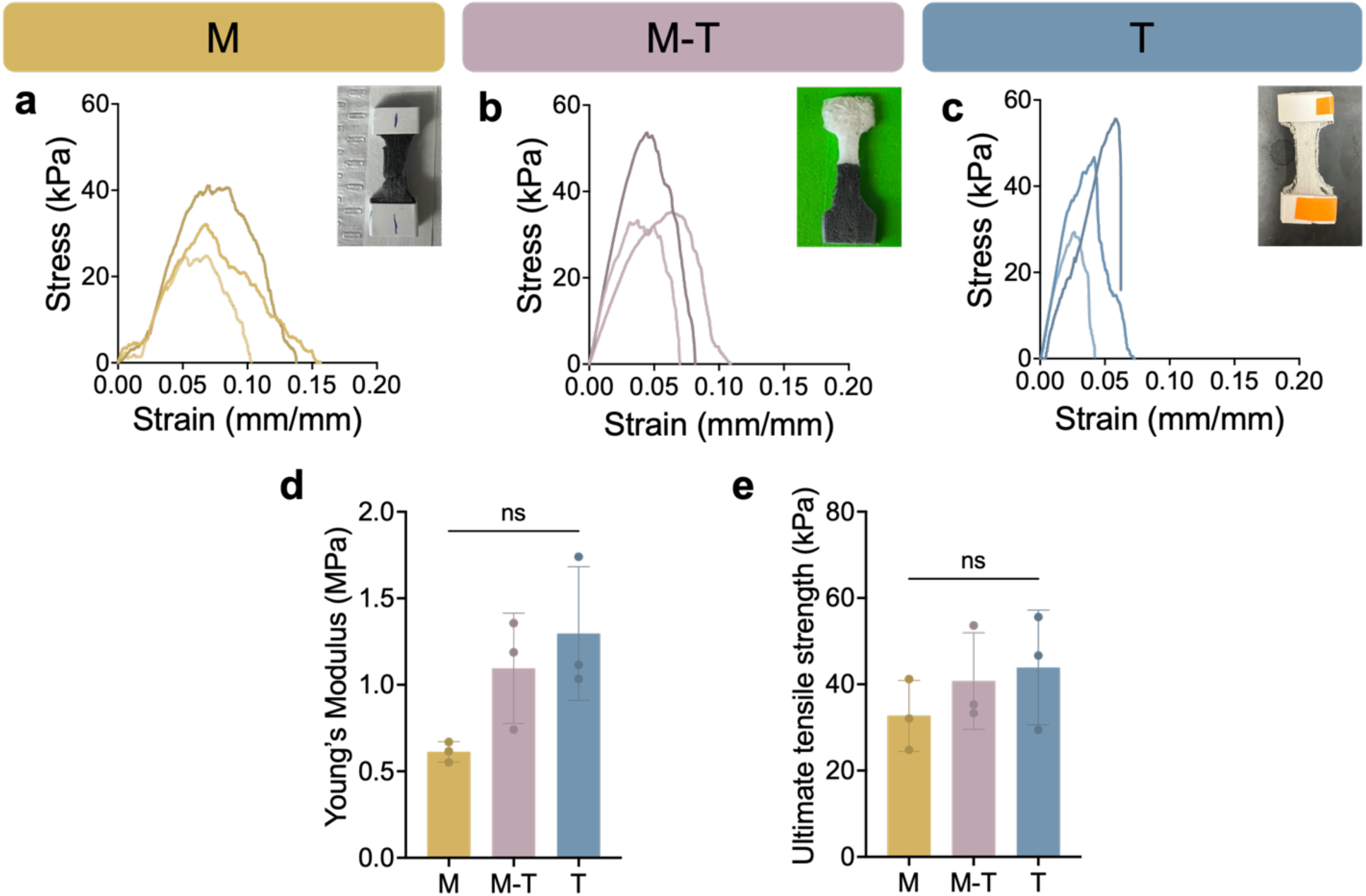
Integrating non-conductive and conductive compartments into a continuous multicompartment scaffold does not diminish mechanics. Representative tensile stress-strain curves of a) single compartment ‘muscle’ (M), b) multicompartment MTJ (M-T), and c) single compartment ‘tendon’ (T) scaffolds under dry conditions. d) Quantification of Young’s modulus and e) ultimate tensile strength reveals comparable mechanical properties across scaffold groups, indicating successful mechanical integration in the multicompartment group. One-way ANOVA with Tukey’s HSD post hoc tests, ns: no statistically significant differences. *N* = 3 scaffolds per experimental group.

Further analysis demonstrated comparable Young’s moduli and ultimate tensile strength values for all scaffold groups (**Figure 4d-e**), indicating successful mechanical integration of the M and T compartments in the multicompartment scaffold. Importantly, the graded scaffold architecture established a mechanically continuous interface between compartments rather than a sharply mismatched junction susceptible to stress concentrations and structural failure. The measured Young’s moduli values in the dry state (∼ 1 MPa) are estimated to correspond to hydrated moduli of ∼ 10 kPa based on previous work with CG scaffolds^[45]^. This stiffness range lies within physiologically relevant skeletal muscle tissue mechanics capable of supporting cellular activity while maintaining structural stability. While some reports indicate that conductive polymer incorporation can compromise scaffold mechanical performance due to the brittle nature of their rigid conjugated backbones^[52]^, we did not observe this phenomenon, potentially due to incorporation of the conductive polymers as distributed particles rather than direct polymerization within the scaffold backbone. Collectively, these results highlight that non-conductive ‘tendon’ and conductive ‘muscle’ scaffold compartments can be successfully mechanically integrated, fulfilling a key requirement for musculotendinous tissue engineering.

### 3.5 Multicompartment scaffolds support stratified myoblast and fibroblast organization as well as sustained metabolic activity of both cell types

After characterizing the promising structural and mechanical properties of the multicompartment scaffold for MTJ tissue engineering, we next evaluated the behavior of seeded myoblasts and fibroblasts. C2C12 myoblasts were seeded within the M compartment, NIH 3T3 fibroblasts within the T compartment, and a combination of both cell types at the M-T interface **(Figure 5a)**. Cell-laden scaffolds were cultured in myoblast growth media (GM) for 4 days, followed by differentiation media (DM) for 4 days to promote myogenic differentiation **(Figure 5b)**. Cell metabolic activity was quantified using a non-destructive alamarBlue assay, with fluorescence intensity serving as an indicator for cell metabolic activity **(Figure 5c)**. Myoblasts exhibited a significant increase in cell metabolic activity from day 1 to day 4, indicating favorable cytocompatibility of the multicompartment scaffold, specifically the PEDOT-containing M compartment. No further significant increase was observed between days 4 and 8, consistent with myogenic differentiation and reduced proliferative activity. A similar trend was observed for fibroblasts in the T compartment, demonstrating sustained metabolic activity even when cultured under myogenic media conditions.

**Figure 5.**
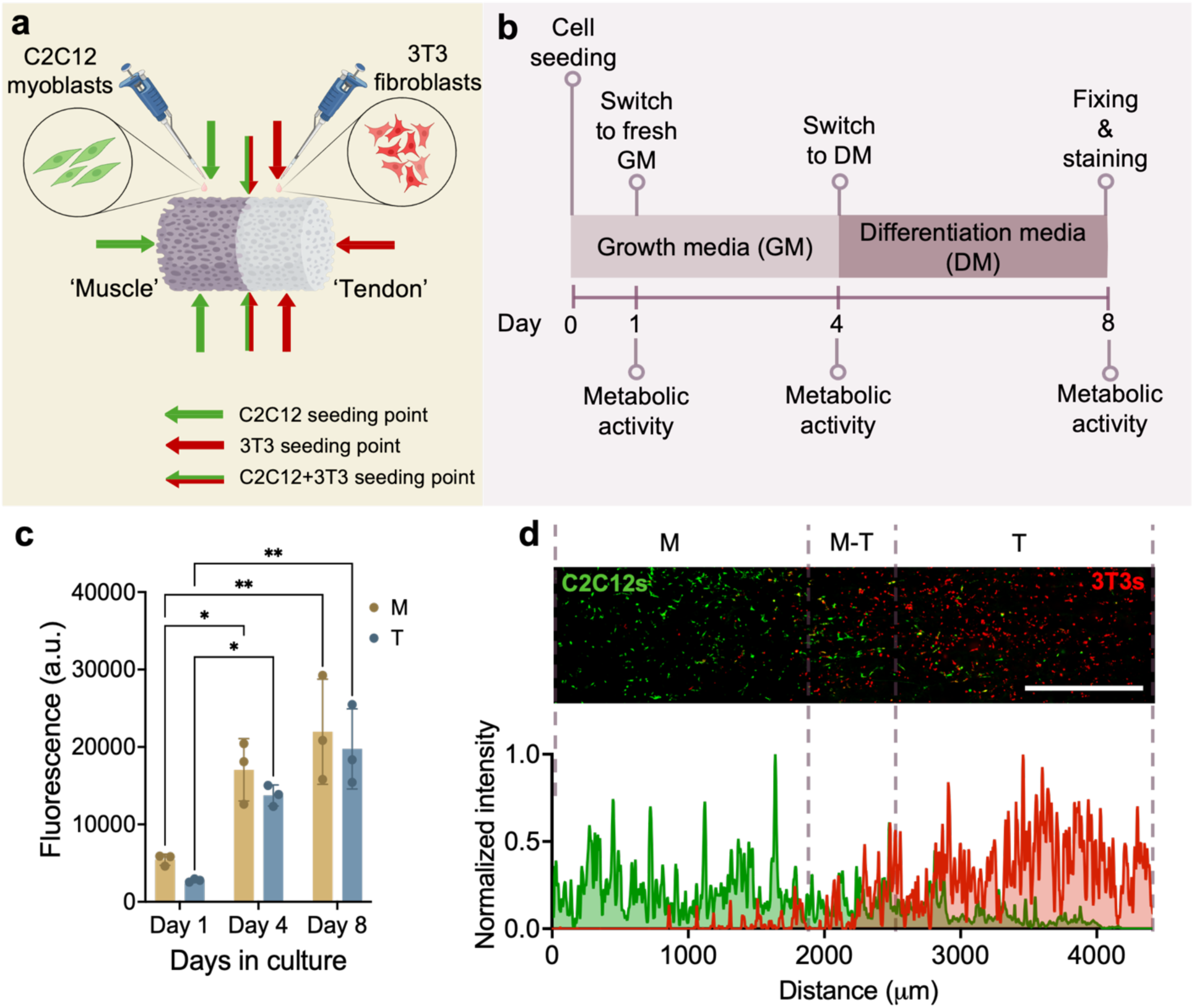
Multicompartment scaffolds support stratified myoblast and fibroblast organization as well as sustained metabolic activity of both cell types. a) Schematic of spatially controlled cell seeding, with C2C12 myoblasts introduced into the ‘muscle’ compartment, NIH 3T3 fibroblasts into the ‘tendon’ compartment, and both cell types present at the interface. b) Experimental timeline illustrating cell seeding, media transitions from growth medium (GM) to differentiation medium (DM), and metabolic activity measurements over an 8-day culture period. c) Quantification of cellular metabolic activity using the alamarBlue assay reveals a time-dependent increase in fluorescence intensity (metabolic activity) in both scaffold compartments. d) Representative fluorescence images and corresponding intensity profiles demonstrating compartment-specific localization of myoblasts (green) and fibroblasts (red), with a mix of both cell populations across the interface region. Two-way ANOVA with Tukey’s HSD post hoc tests, ✽: p < 0.05, ✽✽: p < 0.01. Scale bar: 1000 µm. *N* = 3 scaffolds per experimental group.

Spatial organization of cell populations was assessed via confocal microscopy using cell tracker-labeled myoblasts (green) and fibroblasts (red) **(Figure 5d)**. Distinct compartmental localization of each cell type was observed with myoblasts predominating in the M region and fibroblasts in the T region. Quantitative of fluorescence across the scaffold length further revealed a mix of fluorescent signals corresponding to both myoblasts and fibroblasts across the M-T region. This spatial gradient in cell distribution highlights the ability of the scaffold to support region-specific cellular organization, a critical feature for recapitulating the native MTJ.

### 3.6 Multicompartment scaffolds promote both myoblast and fibroblast alignment

Native skeletal muscle exhibits highly aligned muscle fiber architecture directed by ECM organization, which is essential for coordinated force transmission and tissue function^[53,54]^. Disruption of this structural alignment is associated with impaired muscle function and certain muscle pathologies^[55]^. Biomaterial systems providing aligned structural cues have been shown to enhance myoblast alignment and promote myogenic differentiation^[56]^. Similarly, tendon tissue consists of hierarchically aligned collagen fibers, and instructive biomaterial environments providing anisotropic structural cues have shown to direct cellular alignment and promote tenogenic differentiation of human tendon stem/progenitor cells^[57,58]^.

To evaluate cellular organization within the multicompartment scaffolds, cell cytoskeletal alignment of C2C12 myoblasts and NIH 3T3 fibroblasts was assessed through confocal imaging of F-actin organization together with scaffold backbone morphology (**Figure 6a-c**). Both cell populations exhibited pronounced alignment along the longitudinal scaffold axis, consistent with anisotropic pore architecture generated through directional freeze-drying. Quantitative orientation analysis further confirmed this behavior, with polar plots demonstrating strong frequency accumulation around 0°, the direction of heat transfer during freeze-drying (**Figure 6d**). Importantly, scaffold backbone orientation exhibited corresponding alignment profiles that closely mirrored cytoskeletal organization (**Figure 6e**). These findings confirm that the aligned pore architecture provides robust contact guidance cues capable of directing cellular organization independent of scaffold compartment.

**Figure 6.**
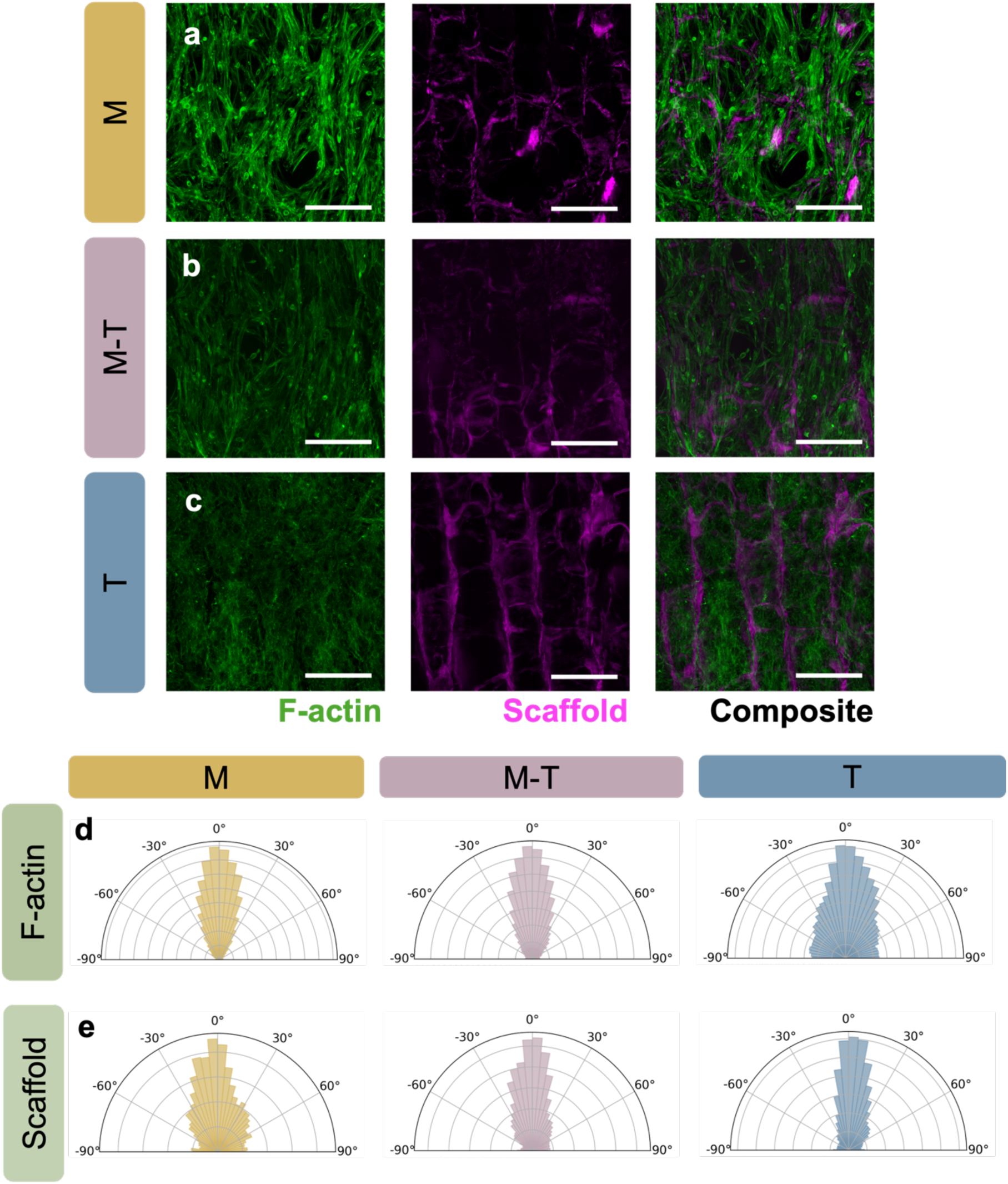
Multicompartment scaffolds promote both myoblast and fibroblast alignment. a-c) Representative confocal fluorescence images of the ‘muscle’ (M), ‘muscle-tendon junction’ (M-T), and ‘tendon’ (T) regions. Cells exhibit elongated morphologies aligned along the scaffold’s longitudinal axis across all compartments, consistent with the underlying anisotropic pore structure. Scale bars: 200 μm. d) Polar histograms of F-actin orientation demonstrate a strong alignment along the longitudinal axis in all regions. e) Corresponding analysis of scaffold backbone orientation shows a similar directional distribution, indicating that cellular alignment closely follows scaffold architecture. Together, these results confirm that aligned scaffolds promote uniform cytoskeletal organization independent of scaffold compartment. *N* = 3 scaffolds per group.

### 3.7 Myogenic differentiation culture supports compartment-specific myotube formation while maintaining fibroblast viability

To evaluate region-specific cellular responses within the multicompartment scaffold, cell-seeded scaffolds were cultured under myogenic differentiation conditions and analyzed for myotube formation and overall viability (nuclei count). Myogenic differentiation was assessed through quantification of myosin heavy chain (MHC)^+^ area, myotube length, myotube width, fusion index, and number of nuclei within MHC^+^ cells. Representative confocal images from the M, M-T, and T regions are shown in **Figure 7a-c**. Extensive MHC expression and multinucleated myotube formation were observed within the M compartment, whereas the T compartment exhibited minimal MHC staining, consistent with the predominance of fibroblasts and the absence of myoblasts in this region.

**Figure 7.**
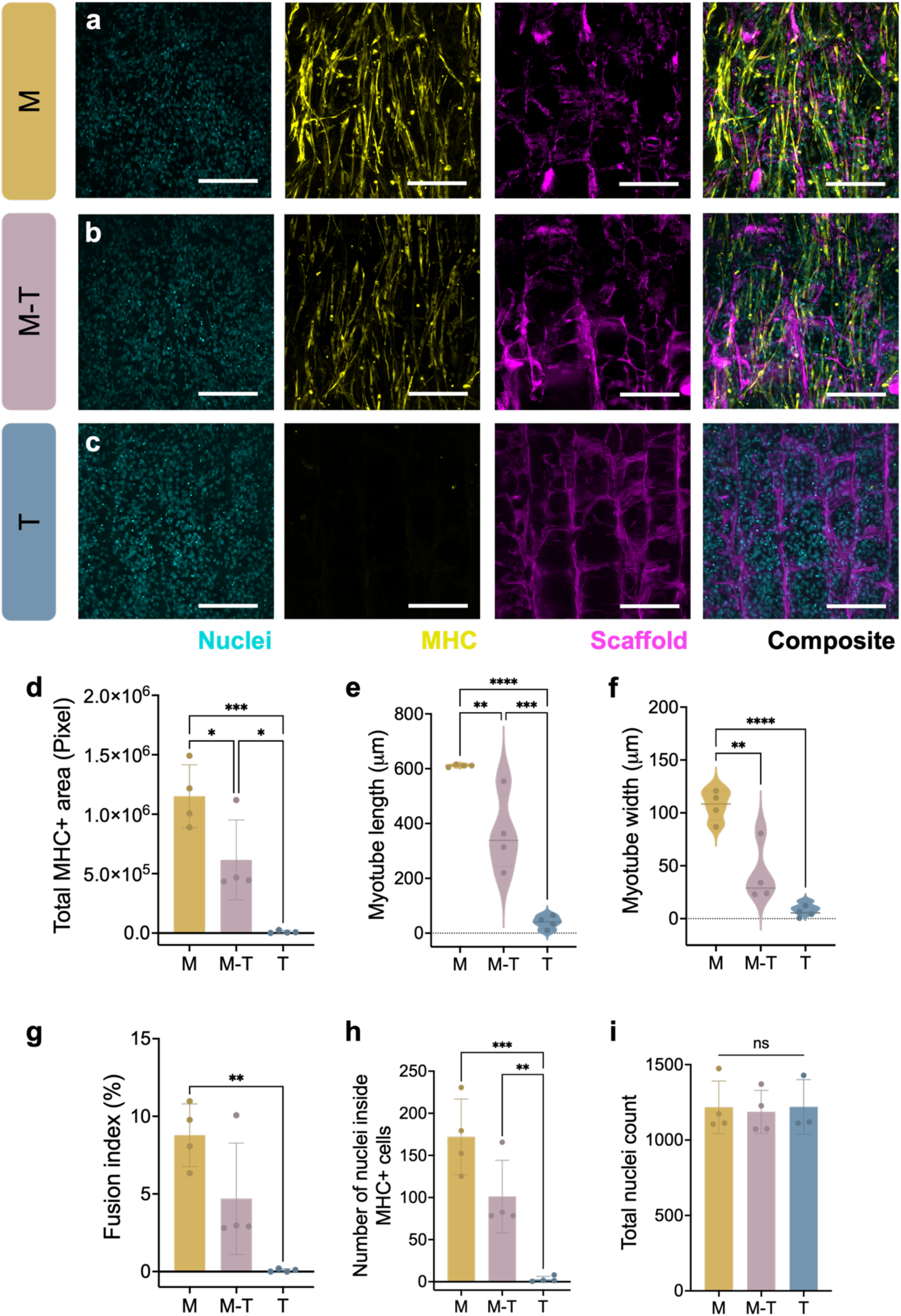
Myogenic differentiation culture supports compartment-specific myotube formation while maintaining fibroblast viability. a-c) Representative fluorescence images of the ‘muscle’ (M), ‘muscle-tendon junction’ (M-T), and ‘tendon’ (T) compartments. Robust myotube formation (MHC⁺ structures) is observed in the M compartment in contrast to reduced differentiation at the interface and minimal to no myotube formation in the T region. Scale bars: 200 μm. d) Quantification of total MHC⁺ area confirms significantly higher myogenic differentiation in the M compartment compared to M-T and T. e, f) Myotube length and width are significantly greater in the M region, with intermediate values at the interface and minimal myotube development in the T compartment. g) Fusion index demonstrates enhanced myoblast fusion in the M compartment relative to the T compartment. h) The number of nuclei within myotubes follows a similar trend, indicating greater maturation in the M compartment. i) Total nuclei count remains comparable across all compartments, confirming consistent cell seeding and overall cell viability throughout the multicompartment scaffold. One-way ANOVA with Tukey’s post hoc tests, ✽: p < 0.05, ✽✽: p < 0.01, ✽✽✽: p < 0.001, ✽✽✽✽: p < 0.0001. *N* = 4 scaffolds per group.

Quantitative analysis demonstrated significantly greater MHC^+^ area in the M region relative to both the M-T and T regions (**Figure 7d**), indicating that the conductive PEDOT-incorporated microenvironment strongly supported myogenic maturation. The M-T region exhibited intermediate MHC expression, significantly greater than the T region yet lower than the M compartment. This confirms the coexistence of both myoblast and fibroblast populations within the interfacial zone. Although MHC expression was expected to be absent in the T compartment due to the presence of fibroblasts, a low level of MHC signal was observed. This minimal expression may be from attachment of suspended myoblasts within the T region during culture.

Consistent with these findings, myotube length and width were significantly increased within the M compartment compared to M-T and T regions (**Figure 7e-f**). Interestingly, myotube lengths within the M-T region displayed a broader distribution, suggesting a gradual transition between myogenic to fibroblastic microenvironments that may recapitulate aspects of the native MTJ interface architecture. Fusion index, calculated as a percentage of nuclei contained within multinucleated MHC^+^ myotubes relative to the total nuclei count, was also significantly elevated in the M compartment (**Figure 7g**), further demonstrating enhanced myogenic differentiation within the conductive scaffold region. The number of nuclei contained within MHC^+^ cells gradually decreased from the M compartment through the M-T region toward the T compartment (**Figure 7h**), further supporting the establishment of a cellular microenvironment across the multicompartment scaffold. Full thickness scaffold confocal imaging revealed globally aligned myotube formation throughout the M compartment, demonstrating that scaffold-induced anisotropic guidance was maintained across entire scaffold thickness rather than limited to localized regions (**Figure S4**).

Importantly, total nuclei count remained comparable across all regions (**Figure 7i**), indicating that fibroblasts retained viability despite exposure to myogenic differentiation media. Collectively, these findings demonstrate that the multicompartment scaffold supports spatially organized myogenic differentiation while simultaneously maintaining fibroblast viability, thereby establishing a permissive microenvironment for formation of a biomimetic musculotendinous junction.

## 4. Conclusions

We developed an aligned multicompartment collagen scaffold incorporating graded structural and biochemical cues as a new tissue engineering platform for the MTJ. Directional freeze-drying of layered conductive and non-conductive collagen suspensions generated a longitudinally aligned porous architecture throughout the scaffold while establishing compartment-specific conductive PEDOT distribution to mimic muscle electrical excitability. Importantly, the layered freeze-drying fabrication approach preserved mechanical continuity throughout the scaffold and prevented abrupt mechanical mismatch at the M-T interface, resulting in a structurally integrated construct containing an interfacial zone similar to the native MTJ. The anisotropic scaffold architecture further provided robust contact guidance cues that directed aligned cellular organization across scaffold compartments. Biologically, the conductive M compartment supported aligned multinucleated myotube formation and enhanced myogenic differentiation, while fibroblasts within the T compartment maintained viability under myogenic differentiation conditions. The M-T region supported coexistence of both cell populations, establishing a graded cellular microenvironment that recapitulates key features of native MTJ. Collectively, this work introduces a multicompartment scaffold platform integrating anisotropic architecture, graded mechanical properties, and spatially controlled electrical conductivity, a previously unexplored parameter in MTJ engineering, to generate a biomimetic *in vitro* MTJ construct. These findings establish a foundation for future studies investigating MTJ-specific cellular interactions and *in vivo* musculotendinous tissue engineering.

## Acknowledgments

The authors thank James Gentry for developing the CellPose code to analyze confocal images, Prof. Geoff Geise for use of his parallel plate cell, and Prof. Silvia Blemker for use of her Instron. SEM images were acquired at the University of Virginia Nanoscale Materials Characterization Facility (NMCF). Some figures were created using BioRender. This work was supported by the NIH (R01AR078866). The content is solely the responsibility of the authors and does not necessarily represent the official views of the National Institutes of Health.

## Data availability statement

All data are available from the authors upon reasonable request.

## References

[1] P. J. Yang, J. S. Temenoff, Tissue Engineering Part B: Reviews 2009, 15, 127.

[2] S. S. Steltzer, A. C. Abraham, M. L. Killian, Curr Osteoporos Rep 2024, 22, 290.

[3] T. Lei, T. Zhang, W. Ju, X. Chen, B. C. Heng, W. Shen, Z. Yin, Bioactive Materials 2021, 6, 2491.

[4] E. Bayrak, P. Yilgor Huri, Front. Mater. 2018, 5, 24.

[5] E. Bayrak, B. Ozcan, J Tissue Sci Eng 2016, 7, DOI 10.4172/2157-7552.1000174.

[6] S. Tong, Y. Sun, B. Kuang, M. Wang, Z. Chen, W. Zhang, J. Chen, Biomedicines 2024, 12, 423.

[7] M. Volpi, A. Paradiso, E. Walejewska, C. Gargioli, M. Costantini, W. Swieszkowski, Adv Healthcare Materials 2024, 13, 2402075.

[8] J. R. Jakobsen, M. R. Krogsgaard, Front. Physiol. 2021, 12, 635561.

[9] M. C. P. Vila Pouca, M. P. L. Parente, R. M. N. Jorge, J. A. Ashton-Miller, Orthopaedic Journal of Sports Medicine 2021, 9, 23259671211020731.

[10] J. W. Prescott, J. S. Yu, American Journal of Roentgenology 2012, 199, W294.

[11] C. P. Dolan, A. R. Clark, J. M. Motherwell, N. B. Janakiram, M. S. Valerio, C. L. Dearth, S. M. Goldman, npj Regen Med 2022, 7, 59.

[12] K. Downing, R. Prisby, V. Varanasi, J. Zhou, Z. Pan, M. Brotto, Current Opinion in Pharmacology 2021, 59, 61.

[13] C. W. Cai, J. A. Grey, D. Hubmacher, W. M. Han, Advances in Wound Care 2024, wound.2024.0079.

[14] F. Snow, C. O’Connell, P. Yang, M. Kita, E. Pirogova, R. J. Williams, R. M. I. Kapsa, A. Quigley, APL Bioengineering 2024, 8, 021505.

[15] T. Y. Kostrominova, S. Calve, E. M. Arruda, L. M. Larkin, 2009.

[16] Y. J. No, M. Castilho, Y. Ramaswamy, H. Zreiqat, Advanced Materials 2020, 32, 1904511.

[17] L. S. G. Bs, Z. G. D. Bs, C. Mora-Navarro, M. B. Fisher, D. O. Freytes, n.d.

[18] B. Charvet, F. Ruggiero, D. L. Guellec, n.d.

[19] A. T. Adams, Z. G. Davis, K. F. Browder, C. L. Dearth, S. M. Goldman, Front. Physiol. 2025, 16, 1555199.

[20] R. J. Monti, R. R. Roy, J. A. Hodgson, V. Reggie Edgerton, Journal of Biomechanics 1999, 32, 371.

[21] N. Narayanan, S. Calve, Connective Tissue Research 2021, 62, 53.

[22] H. H. Lu, S. Thomopoulos, Annual Review of Biomedical Engineering 2013, 15, 201.

[23] A. B. Knudsen, M. Larsen, A. L. Mackey, M. Hjort, K. K. Hansen, K. Qvortrup, M. Kjær, M. R. Krogsgaard, Scandinavian Journal of Medicine & Science in Sports 2015, 25, e116.

[24] J. G. Tidball, G. Salem, R. Zernicke, Journal of Applied Physiology 1993, 74, 1280.

[25] M. Kääriäinen, T. Järvinen, M. Järvinen, J. Rantanen, H. Kalimo, Scandinavian Med Sci Sports 2000, 10, 332.

[26] M. R. Ladd, S. J. Lee, J. D. Stitzel, A. Atala, J. J. Yoo, Biomaterials 2011, 32, 1549.

[27] T. K. Merceron, M. Burt, Y.-J. Seol, H.-W. Kang, S. J. Lee, J. J. Yoo, A. Atala, Biofabrication 2015, 7, 035003.

[28] W. Kiratitanaporn, J. Guan, M. Tang, Y. Xiang, T. Lu, A. Balayan, A. Lao, D. B. Berry, S. Chen, Biomater. Sci. 2024, 12, 6047.

[29] W. J. Kim, G. H. Kim, Bioengineering & Transla Med 2022, 7, e10321.

[30] F. De Paolis, M. Volpi, C. Fuoco, A. Reggio, R. Deodati, S. Bernardini, A. Palma, U. G. Longo, L. Santorelli, F. Scirocchi, et al., Biofabrication 2026, 18, 015035.

[31] N. Alegret, A. Dominguez-Alfaro, D. Mecerreyes, Biomacromolecules 2019, 20, 73.

[32] B. Guo, P. X. Ma, Biomacromolecules 2018, 19, 1764.

[33] Y. Du, J. Ge, Y. Li, P. X. Ma, B. Lei, Biomaterials 2018, 157, 40.

[34] A. F. Quigley, J. M. Razal, M. Kita, R. Jalili, A. Gelmi, A. Penington, R. Ovalle-Robles, R. H. Baughman, G. M. Clark, G. G. Wallace, et al., Advanced Healthcare Materials 2012, 1, 801.

[35] A. Saberi, F. Jabbari, P. Zarrintaj, M. R. Saeb, M. Mozafari, Biomolecules 2019, 9, 448.

[36] R. Balint, N. J. Cassidy, S. H. Cartmell, Acta Biomaterialia 2014, 10, 2341.

[37] I. M. Basurto, M. T. Mora, G. M. Gardner, G. J. Christ, S. R. Caliari, Biomater. Sci. 2021, 9, 4040.

[38] I. M. Basurto, S. A. Muhammad, G. M. Gardner, G. J. Christ, S. R. Caliari, J Biomedical Materials Res 2022, jbm.a.37418.

[39] M. Shi, R. Dong, J. Hu, B. Guo, Chemical Engineering Journal 2023, 457, 141110.

[40] Kenry, B. Liu, Biomacromolecules 2018, 19, 1783.

[41] L. Ghasemi-Mobarakeh, M. P. Prabhakaran, M. Morshed, M. H. Nasr-Esfahani, H. Baharvand, S. Kiani, S. S. Al-Deyab, S. Ramakrishna, J Tissue Eng Regen Med 2011, 5, e17.

[42] S. R. Caliari, B. A. C. Harley, Biomaterials 2011, 32, 5330.

[43] G. C. Bandara, R. D. Boudreau, W. Wyatt, S. R. Caliari, J Biomedical Materials Res 2026, 114, e70048.

[44] S. R. Caliari, D. W. Weisgerber, W. K. Grier, Z. Mahmassani, M. D. Boppart, B. A. C. Harley, Adv. Healthcare Mater. 2015, 4, 831.

[45] B. Harley, J. Leung, E. Silva, L. Gibson, Acta Biomaterialia 2007, 3, 463.

[46] L. H. H. Olde Damink, P. J. Dijkstra, M. J. A. Van Luyn, P. B. Van Wachem, P. Nieuwenhuis, J. Feijen, Biomaterials 1996, 17, 765.

[47] S. N. Rampersad, Sensors 2012, 12, 12347.

[48] J. L. Gentry, S. R. Caliari, n.d.

[49] K. J. Cutler, C. Stringer, T. W. Lo, L. Rappez, N. Stroustrup, S. Brook Peterson, P. A. Wiggins, J. D. Mougous, Nat Methods 2022, 19, 1438.

[50] D. T. Tran, L. Tsai, J Biol Phys 2023, 49, 257.

[51] C. N. Maganaris, P. Chatzistergos, N. D. Reeves, M. V. Narici, Front. Physiol. 2017, 8, DOI 10.3389/fphys.2017.00091.

[52] B. Guo, Z. Ma, L. Pan, Y. Shi, J Polym Sci B Polym Phys 2019, 57, 1606.

[53] R. P. Wohlgemuth, S. E. Brashear, L. R. Smith, American Journal of Physiology-Cell Physiology 2023, 325, C1017.

[54] A. R. Gillies, R. L. Lieber, Muscle Nerve 2011, 44, 318.

[55] K.-Y. Lee, H.-X. Loh, A. Wan, Micromachines 2021, 13, 71.

[56] S. L. Hume, S. M. Hoyt, J. S. Walker, B. V. Sridhar, J. F. Ashley, C. N. Bowman, S. J. Bryant, Acta Biomaterialia 2012, 8, 2193.

[57] Z. Yin, X. Chen, J. L. Chen, W. L. Shen, T. M. Hieu Nguyen, L. Gao, H. W. Ouyang, Biomaterials 2010, 31, 2163.

[58] S. Testa, M. Costantini, E. Fornetti, S. Bernardini, M. Trombetta, D. Seliktar, S. Cannata, A. Rainer, C. Gargioli, J Cellular Molecular Medi 2017, 21, 2711.

