## Supplemental Information for "Aligned multicompartment collagen scaffolds support stratified myoblast and fibroblast behavior for musculotendinous tissue engineering"

| **Scaffold compartment** | **M** | **M-T** | **T** |
|---|---|---|---|
| **Longitudinal pore size (µm)** | 133.1 ± 8.1 | 125.0 ± 7.5 | 129.2 ± 3.5 |
| **Transverse pore size (µm)** | 116.9 ± 3.0 | 119.6 ± 5.5 | 122.0 ± 1.6 |

**Table S1. Quantification of scaffold pore size.** Mean ± standard deviations of longitudinal and transverse pore size quantification are shown for ‘muscle’ (M), ‘muscle-tendon junction’ (M-T), and ‘tendon’ (T) scaffold compartments. *N* = 3 scaffolds per experimental group.

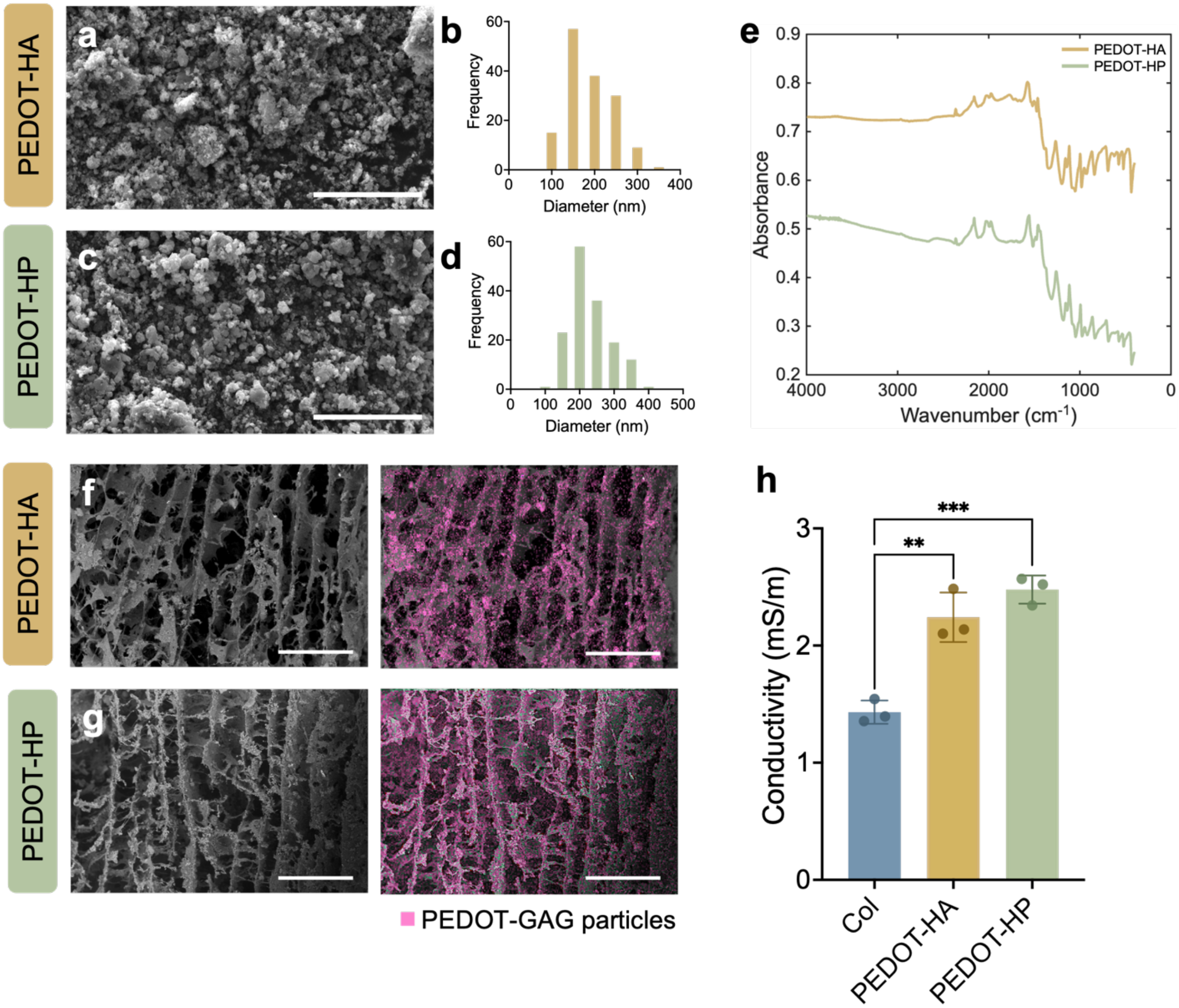


**Figure S1. GAG-doped PEDOT particles enhance scaffold electrical conductivity.** a, c) Scanning electron microscopy (SEM) images of PEDOT-HA and PEDOT-HP particles. b, d) Corresponding particle size distributions indicate comparable size ranges with mean diameters of 190 ± 55 nm for PEDOT-HA and 229 ± 56 nm for PEDOT-HP. e) Fourier transform infrared (FTIR) spectra confirm successful incorporation of GAG dopants within PEDOT. Characteristic PEDOT absorption bands corresponding to thiophene ring C=C stretching (1550-1570 $cm^{-1}$), C-O stretching (1100-1260 $cm^{-1}$), and C-S vibrations (700-990 $cm^{-1}$) were observed in both PEDOT-HA and PEDOT-HP, confirming successful polymerization of EDOT. Additional sulfate- (1430 $cm^{-1}$) and carboxylate- (1760 $cm^{-1}$) associated bands further verified incorporation of HP and HA respectively. f, g) Representative SEM images of scaffolds incorporating PEDOT-HA and PEDOT-HP particles respectively, with corresponding energy-dispersive X-ray spectroscopy (EDS) overlays highlighting the spatial distribution of PEDOT-GAG particles (pink), demonstrating uniform incorporation within the scaffolds. h) Electrical conductivity measurements, obtained via linear sweep voltammetry, show a significant increase in conductivity for PEDOT-HA and PEDOT-HP scaffolds compared to collagen-only controls (Col). One-way ANOVA with Tukey's post hoc test. ✻✻: $p < 0.01$, ✻✻✻: $p < 0.001$. Scale bars: 15 μm (a, b), 300 μm (f, g); $N$ = 3 scaffolds per experimental group.

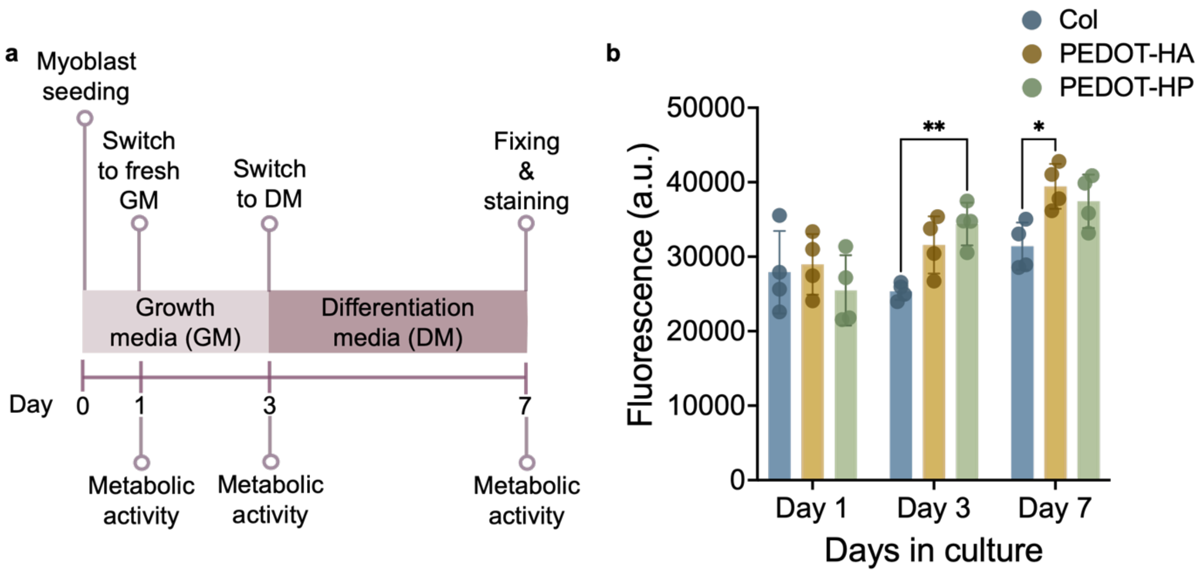


**Figure S2. Scaffolds incorporating GAG-doped PEDOT particles support sustained myoblast metabolic activity.** a) Schematic illustrating the experimental timeline for cell seeding, media transitions from growth media (GM) to differentiation media (DM), and metabolic activity measurements over a 7-day culture period. b) Quantification of myoblast metabolic activity using the alamarBlue assay demonstrates sustained metabolic activity in all experimental groups, highlighting the cytocompatibility of the GAG-doped PEDOT particles. Two-way ANOVA with Tukey's HSD post hoc tests. ✻: $p < 0.05$, ✻✻: $p < 0.01$. $N = 4$ scaffolds per experimental group.

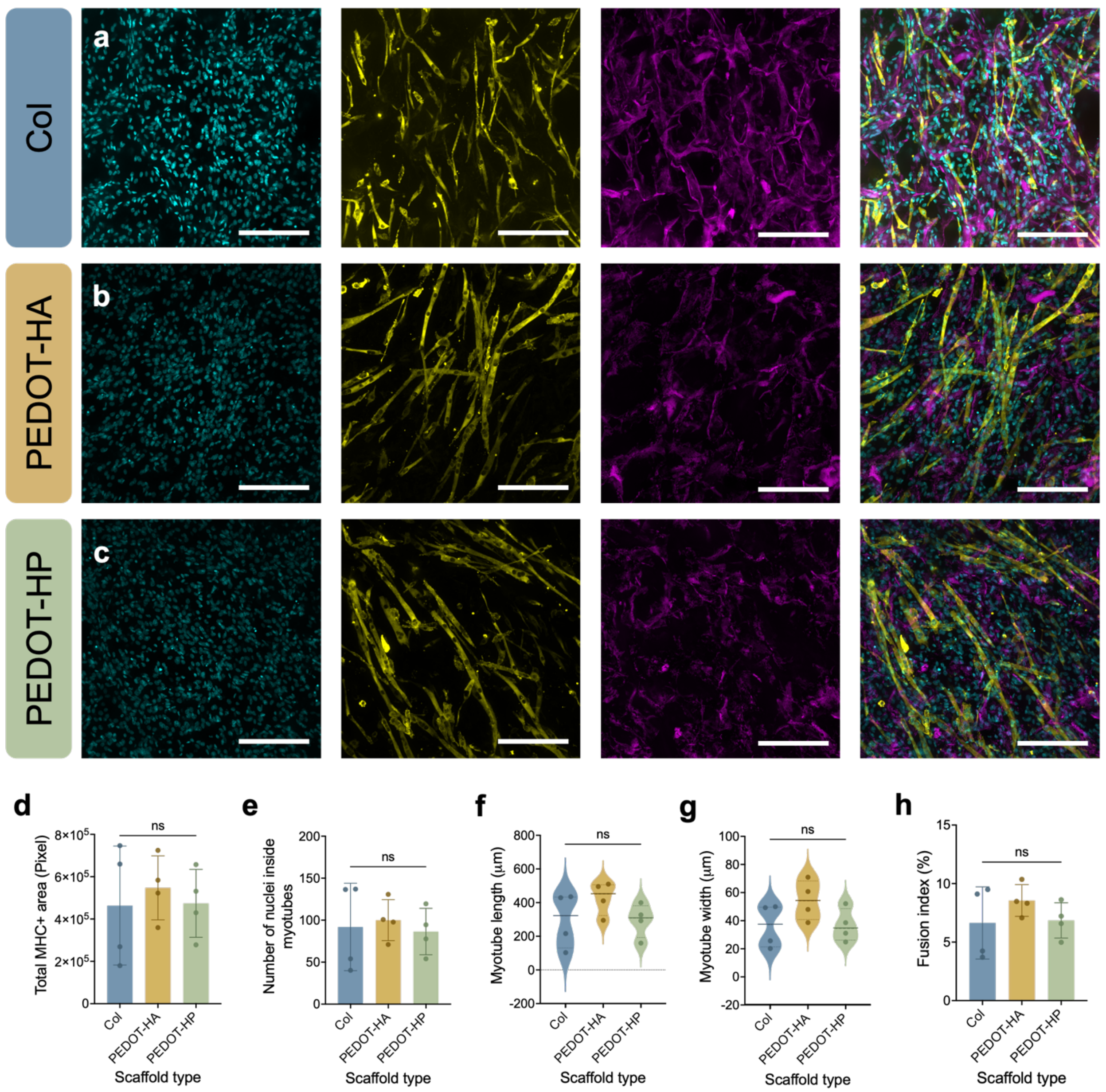


**Figure S3. Scaffolds incorporating GAG-doped PEDOT particles support myogenic differentiation after 4 days in differentiation conditions.** a-c) Representative fluorescence images of collagen-only (Col), PEDOT-HA, and PEDOT-HP particle-laden scaffolds. Scale bars: 200 µm. All groups support similar levels of myotube formation as measured by d) quantification of total MHC$^+$ area, e) number of nuclei within myotubes, f) myotube length, g) myotube width, and h) fusion index. However, the PEDOT-HA scaffolds support the highest levels of myogenic differentiation by all measures. One-way ANOVA with Tukey's HSD post hoc tests, ns: no statistically significant differences. *N* = 4 scaffolds per experimental group.

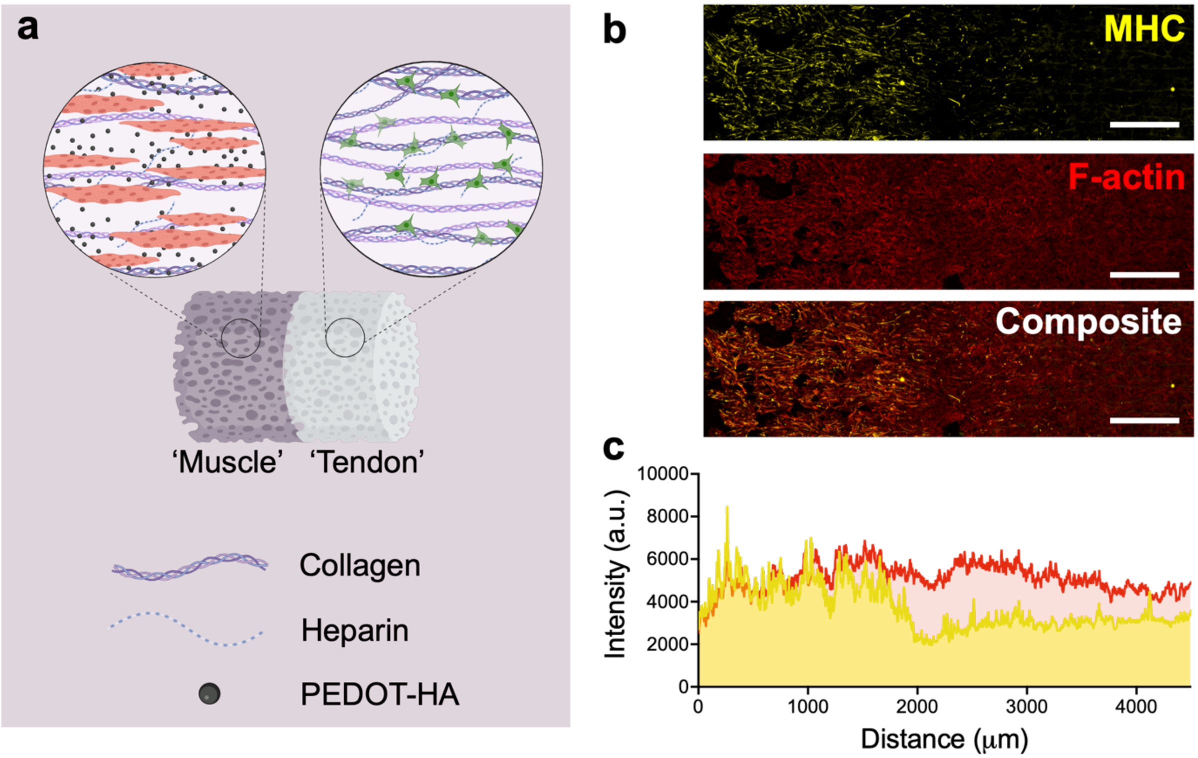


**Figure S4. Multicompartment scaffold architecture supports stratified myotube maturation.** a) Schematic representation of compartment-specific scaffold composition and corresponding cellular organization within the 'muscle' (M) and 'tendon' (T) regions following culture under myogenic differentiation conditions. b) Representative confocal images of myosin heavy chain (MHC), F-actin, and composite overlays demonstrate myotube organization primarily in the muscle compartment and aligned F-actin expression throughout the entire scaffold. c) Spatial intensity profiles of MHC and F-actin expression across the scaffold further demonstrate consistency of the F-actin signal and a transition in MHC expression at the muscle-tendon compartment junction. Scale bars: 1000 μm.
